# Prefrontal representation of affective stimuli: importance of stress, sex, and context

**DOI:** 10.1101/2022.09.02.506435

**Authors:** Tyler Wallace, Brent Myers

**Affiliations:** Biomedical Sciences, Colorado State University, Fort Collins, CO

**Author notes:** **Address for correspondence** Brent Myers, Ph.D., Department of Biomedical Sciences, Colorado State University, 1617 Campus Delivery, Fort Collins, CO 80523.

**Keywords:** infralimbic cortex, chronic variable stress, social, food reward, photometry, ovariectomy

## Abstract

Major depressive disorder diminishes health-related quality-of-life and disproportionality impacts females. Negative life events predict the onset of depressive symptoms including negative mood, anhedonia, and social withdrawal. While the neurobiology of affective illness is not completely understood, structural and functional changes in the ventromedial prefrontal cortex (vmPFC) associate with mood and anxiety disorders. Rodent studies have investigated the prefrontal impact of chronic stress, a preclinical model of mood disruption. However, *ex vivo* slice and postmortem histological studies have focused largely on males and yielded mixed results. Although genetically-defined recordings in behaving animals of both sexes have not been reported. Here, we hypothesized that chronic variable stress (CVS) would reduce the neural processing of affective stimuli in the infralimbic region of rodent vmPFC region. To test this hypothesis, we targeted expression of a fluorescent calcium indicator, GCaMP6s, to infralimbic pyramidal cells. In males, CVS reduced infralimbic responses to social interaction and restraint stress but increased responses to novel objects and food reward. In contrast, females did not have CVS-induced changes in infralimbic activity, which was partially dependent on ovarian status. Collectively, these results indicate that both male and female vmPFC pyramidal cells encode social, stress, and reward stimuli. However, chronic stress effects are sex-dependent and behavior-specific. Ultimately, these findings extend the understanding of chronic stress-induced prefrontal dysfunction and indicate that sex is a critical factor for cortical processing of affective stimuli.

**Significance:** Mood disorders are often preceded by stressful experiences and exhibit sex differences in prevalence. Further, affective illness is frequently accompanied by changes in prefrontal cortical function. However, the neurobiology that underlies sex-specific cortical processing of affective stimuli is poorly understood. Therefore, the current studies employ cell type-specific calcium photometry to determine the effects of chronic stress on ventromedial prefrontal cortex (vmPFC) activity in male and female rats. Our results indicate that chronic stress exposure differentially alters vmPFC encoding of social, stress, and reward stimuli. Moreover, the sex-specific effects of chronic stress are partially mediated by ovarian hormones. Ultimately, our results indicate that prolonged stress alters vmPFC function in a context-dependent and sex-specific manner, contributing to the biological bases for sex differences in emotional processing.

## Introduction

Major depressive disorder (MDD), characterized by negative mood, anhedonia, and avoidance behaviors, is the leading cause of years lived with disability worldwide (Friedrich, 2017). Furthermore, females are disproportionately impacted by depressive disorders, with twice the prevalence of males (Kuehner, 2017). Epidemiological studies implicate life stressors as a primary risk factor for the onset of mood disorders (Grippo and Johnson, 2009; Binder and Nemeroff, 2010; Myers et al., 2014; Sgoifo et al., 2015). Additionally, patients experiencing depression have exaggerated physiological responses to novel stressors and impaired habituation to repeated stressors (Gianferante et al., 2014; Morris and Rao, 2014). However, the mechanisms that integrate the impacts of biological sex and stress history on neural processing of affective stimuli are largely unexplored.

The ventromedial prefrontal cortex (vmPFC) is important for cognitive and emotional processes; furthermore, altered structure and function of vmPFC associates with mood disorders (Krishnan and Nestler, 2008; Wallace and Myers, 2021; Wood and Grafman, 2003; Myers-Schulz and Koenigs, 2012; McKlveen et al., 2015). In particular, the Brodmann Area 25 (BA25) subregion of vmPFC is activated by sadness-provoking stimuli, responds to social isolation, and has reduced volume in MDD (Liotti et al., 2000; Beckmann et al., 2009; Vijayakumar et al., 2017). Although, studies of the relationship between BA25 activity and affective illness have yielded mixed results. Combined-sex studies have reported both decreased activity in MDD (Drevets et al., 1997) as well as increased activity in treatment-resistant depression (Mayberg et al., 2005). Thus, while vmPFC function relates to mood, the role of defined cell populations in regulating emotion is unclear.

The rodent infralimbic cortex (IL), a homolog of BA25, contains glutamatergic pyramidal neurons as well as a heterogenous network of interneurons (McKlveen et al., 2015, 2016). Collectively, IL local circuitry regulates activity of the pyramidal cell projections that target the limbic system to modulate behavior and physiology (Vertes, 2004; Wood et al., 2019). In fact, acute optogenetic stimulation of rat IL pyramidal cells regulates socio-affective behaviors and physiological stress responses in a sex-dependent manner (Wallace et al. 2021). Further, chronic stress exposure induces dendritic atrophy of male IL pyramidal cells (Cook and Wellman, 2004; Cerqueira et al., 2005; Goldwater et al., 2009; Shansky et al., 2009; Czéh et al., 2018), an effect prevented in females by ovarian hormones (Shansky et al., 2010; Wei et al., 2014). Although multiple studies provide histological evidence for altered pyramidal morphology, functional studies have yielded conflicting results. To date, *ex vivo* slice recordings have focused on males and found evidence of chronic stress both increasing pyramidal cell inhibition (McKlveen et al., 2016) and decreasing inhibition (Czéh et al., 2018). Combined with reports of chronic stress-induced interneuronal plasticity and altered gene expression (Gilabert-Juan et al., 2013; McKlveen et al., 2016, 2019; Shepard et al., 2016; Czéh et al., 2018; Page et al., 2019), it is unclear how prolonged adversity impacts the endogenous activity of vmPFC pyramidal cells.

To determine how vmPFC neural populations represent affective stimuli *in vivo*, we expressed the fluorescent calcium indicator GCaMP6 under the calcium/calmodulin-dependent protein kinase type II α (CaMKIIα) promoter. Photometric recordings of IL^CaMKIIα^ pyramidal neuron activity were carried out during exposure to novel, social, stress, and reward stimuli in control rats, as well as animals exposed to chronic variable stress (CVS). To test the hypothesis that chronic stress-induced shifts in excitatory/inhibitory balance are sex-specific, experiments included male, female, and ovariectomized (OVX) rats. Altogether, these studies provide evidence that behavioral context, stress history, and biological sex are pivotal factors affecting the real-time activity of IL^CaMKIIα^ cells in behaving animals.

## Methods

### Animals

Age-matched adult male (250-300 g), female (150-200 g), and ovariectomized (OVX) female (250-300 g) Sprague-Dawley rats were obtained from Envigo (Denver, CO). Ovariectomy occurred two weeks before arrival. After stereotaxic surgery, rats were housed individually in shoebox cages with cardboard tubes for enrichment in a temperature- and humidity-controlled room with a 12-hour light-dark cycle (lights on at 07:00 h, off at 19:00 h) and food and water *ad libitum*. Per ARRIVE guidelines, all treatments were randomized, and experimenters were blinded. All procedures and protocols were approved by the Colorado State University Institutional Animal Care and Use Committee (protocol: 2129) and complied with the National Institutes of Health Guidelines for the Care and Use of Laboratory Animals. Signs of poor health and/or weight loss ≥ 20% of pre-surgical weight were a priori exclusion criteria. These criteria were not met by any animals in the current experiments

### Microinjection and cannulation

Microinjections and cannulations were performed as previously described (Wallace et al., 2021). Briefly, rats were anesthetized with isoflurane (1-5%) and administered analgesic (0.6 mg/kg buprenorphine-SR, subcutaneous) and antibiotic (5 mg/kg gentamicin, intramuscular). Rats received unilateral microinjections (1-1.5 µL) of adeno-associated virus (AAV) into the IL (males: 2.7 mm anterior to bregma, 0.6 mm lateral to midline, and 4.2 mm ventral from dura; females: 2.3 mm anterior to bregma, 0.5 mm lateral to midline, and 4 mm ventral from dura) in randomized fashion so that right and left IL were equally represented. AAV9-packaged constructs (Addgene, virus # 107790) expressed GCaMP6s under the CaMKIIα promoter to achieve pyramidal cell-predominant expression (Wood et al., 2019). All microinjections were carried out with a 25-gauge, 2-µL microsyringe (Hamilton, Reno, NV) using a microinjection unit (Kopf, Tujunga, CA) at a rate of 5 min/µL. Following injection, unilateral fiber-optic cannulas (flat tip 400/430 μm, NA = 0.57, 5 mm protrusion; Doric Lenses, Québec, Canada) were aligned with the IL injection sites and lowered to the IL. Cannulas were secured to the skull with metal screws (Plastics One) and dental cement (Stoelting, Wood Dale, IL). Skin was sutured and, following 1 week of recovery, rats were handled daily and acclimated to the recording procedure for another week before experiments began. Rat handling and cannula habituation continued daily throughout experiments.

### Experimental design

Multiple experiments were conducted with results displayed collectively given the normalization of photometry data. Experiment 1a was carried out in No CVS and CVS males (n = 15/group) according to the timeline in **Figure 1B**. Here, responses to social, stress, and reward stimuli were assessed. Due to the possibility of metabolic state affecting IL^CaMKIIα^ responses to food reward, an additional experiment (experiment 1b) was carried out with weight-matched No CVS males (n = 7). Experiment 2 included intact cycling females that were either No CVS controls or exposed to CVS (n = 10/group). A final experiment was run in OVX females to determine whether ovarian hormones modulate IL^CaMKIIα^ neural activity in No CVS controls and CVS-exposed animals (n= 11/group). Experiments 2 and 3 followed the same experimental timeline outlined in **Figure 1B** with tissue collected at the conclusion of all experiments to verify fiber optic placement.

**Figure 1:**
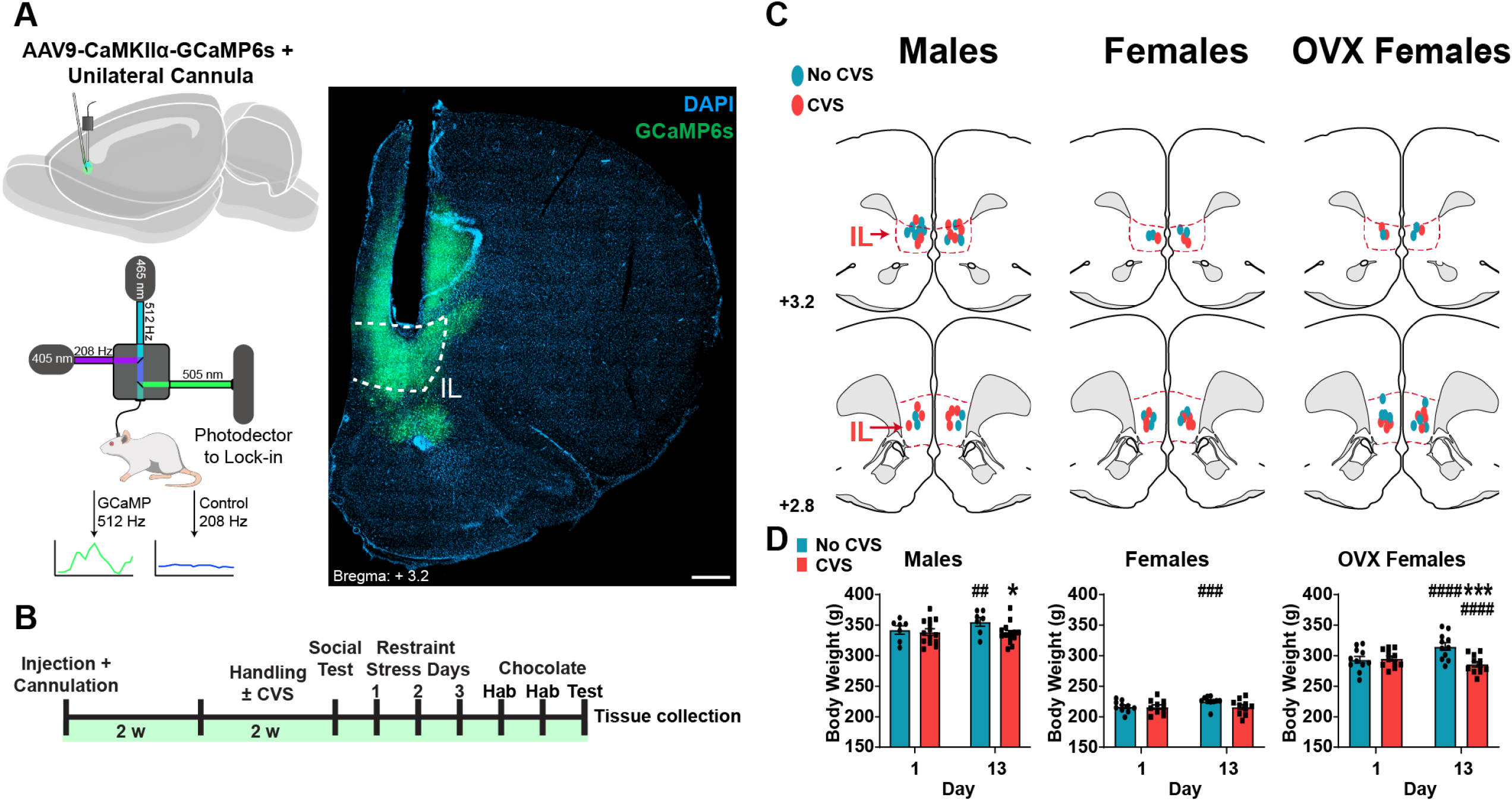
Experimental design. (**A**) Top left: Schematic of AAV injections and cannulation. Bottom left: schematic of fiber photometry recording system. Right: photomicrograph of GCaMP6s expression at the IL cannulation site; scale bar: 500 μm. (**B**) Experimental timeline; CVS: chronic variable stress. (**C**) Mapped fiber optic cannula positions within, or at the dorsal margin of, the IL (red outline); numbers are mm rostral to bregma, OVX: ovariectomized. (**D**) Body weight of animals before and after stress condition, CVS or No CVS. Within-treatment comparisons across time: ^##^ *p <* 0.01, ^###^ *p <* 0.001, ^####^ *p <* 0.0001. Within-time comparisons between treatments: * *p <* 0.05, *** *p <* 0.001.

### CVS

CVS was conducted as previously described (Myers et al., 2017; Pace et al., 2020a; Wallace et al., 2021) with twice daily (AM and PM) heterotypic stressors presented in a randomized manner including: exposure to a brightly-lit open field (28.5 × 18’’, 13’’ deep, 1 hour), cold room (4° C, 1 hour), cage tilt (45°, 1 hr), forced swim (23° to 27° C, 10 min), predator odor (fox or coyote urine, 1 hr), and shaker stress (100 rpm, 1 hr). Additionally, overnight stressors were variably included, comprised of damp bedding (400 mL water) or overnight light. Rats were weighed every 3-4 days to monitor body weight over the course of experimentation.

### Food restriction

A cohort of male rats (experiment 1b) was food restricted for 14 days to control for the reduced food intake and body weight gain of CVS-exposed males. Two weeks after injection and cannulation, male rats received 16 grams of standard chow at lights off and 2 grams of standard chow at lights on to prevent fasting, as previously described (Flak et al., 2011; Schaeuble et al., 2019). Similar to all other experiments, food-restricted rats were given chocolate chips for 2 days prior to test day with food reward testing performed on day 15.

### Estrous cycle

All intact cycling female rats were housed concurrently and followed the same experimental design. Immediately following all tests, vaginal cytology was analyzed to approximate the estrous cycle stage as previously described (Wallace et al., 2021). Briefly, a cotton swab dipped in deionized water was used to collect and transfer cells onto a glass slide. Dried slides were viewed via light microscopy (10x objective) by a minimum of two blind observers to categorize as proestrus, estrus, metestrus, or diestrus (Cora et al., 2015; Solomon et al., 2015).

### Fiber photometry

GCaMP6s fluorescence was recorded to measure activity-dependent changes in intracellular calcium (Gunaydin et al., 2014). Implanted fiber optic cannulas were connected via patch cord to a fiber photometry system (Doric: 1-site 2-color Fiber Photometry System). Excitation was achieved thrfough a 465 nm wavelength LED modulated at 512 Hz. Movement control was achieved with 405 nm wavelength LED modulated at 208 Hz. Power delivery was approximately 3 mW at cannula tip, measured with a photodiode sensor (PM160, Thorlabs Inc, Newton, NJ). To allow for rodent movement, rats were connected to the photometry system through a pigtailed rotary joint patch cord. Emitted fluorescence was captured with a photon detector (Newport Model: 2151). To separate the components of the 465 nm and 405 nm excitation, a lock-in amplifier filtered the incoming fluorescence into the 512 Hz and 208 Hz components respectively with a 12 Hz band filter. Autofluorescence was reduced by running the 465 and 405 nm LEDs at full power for 30 min prior to testing. Further, the optical interface between the implanted fiber optic and the patch cord was cleaned with 10% ethanol and dried immediately before testing.

Photometry recordings were acquired at 1200 Hz and down sampled to 30 Hz for analysis (MATLAB). Behavior was recorded by a camera mounted above the arena for automated optic hardware control (Noldus Information Technologies). A transistor-transistor logic pulse at the beginning of video acquisition was used to timeclock with photometry recordings. Prior to analysis, 465 nm and 405 nm components of the excited 512 nm fluorescence were assessed by a treatment-blind observer for movement artifacts. If movement artifacts were detected, the recorded trials were excluded from the analysis. Photometry signal analysis and z-score calculation were performed using MATLAB (version 2019b, MathWorks).

### Novel object and social interaction

A modified version of a social behavior arena was used to accommodate optic patch cords (Felix-Ortiz and Tye, 2014; Moy et al., 2004). To reduce environmental novelty, experimental rats were habituated to the interaction arena for 3 days (10 min/day) prior to testing. Additionally, interactor rats were habituated to enclosures in the arena for 3 days (20 min/day) prior to testing. On testing day, the 14^th^ day of CVS, rats were connected to fiber optic patch cords and placed in the center of a black rectangular Plexiglas arena (36 × 23’’, 15.8’’ deep). Initially, the arena was empty and experimental rats were allowed to explore. After 5 min, an empty enclosure, novel to the experimental animals, was placed in the center of the arena. Rats were then allowed to explore the novel object for 5 min, after which the empty enclosure was removed. After 2 min, a new enclosure containing a novel conspecific rat was placed in the center of the arena for 5 min. Interactions were scored by a treatment-blind observer as nose pokes onto the empty enclosure (object interaction) or enclosure containing a conspecific (social interaction).

For analysis of IL^CaMKIIα^ GCaMP fluorescence during interactions, the 465 nm component signal was median Z-scored with a rolling 5-min median and median absolute deviation (MAD), 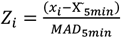, then aligned to interactions and averaged within subjects. The within-subject mean signals were then averaged within groups for statistical analysis. Object interactions were filtered to identify interactions that were ≥ 1 sec and preceded by 3 sec of non-interaction. Social behaviors were filtered to detect interactions ≥ 2 sec and preceded by 3 sec of non-interaction. The first object and social interactions were analyzed separately from subsequent interactions to examine contextual novelty. To determine if the GCaMP signal differed during interactions from non-specific behavior, 7 sec intervals without interactions were randomly selected with the average GCaMP signal calculated during the middle 2 sec of these periods. To isolate IL^CaMKIIα^ activity prior to interaction, social interactions were filtered to only include interactions with a preceding ≥ 10-sec period of non-interaction.

### Restraint stress and endocrine analysis

To examine IL^CaMKIIα^ cellular responses to novel and repeated stress animals underwent 3 days of restraint. To measure baseline neural activity, rats were connected to the photometry system in the home cage for 5 min each day. Rats were then restrained in plastic film decapicones and reconnected to the photometry system for 30 min. To assess endocrine stress responses, blood samples (approximately 250 μL) were collected by tail clip at the end of restraint (Vahl et al., 2005). All samples were collected following the recording session so that neural responses were specific to restraint stress.

For analysis, the 465 nm photometry signal was median Z-scored with a rolling 1-min window within 5-min bins to match the home cage period. Transient peaks in calcium fluorescence were defined as values exceeding 2.91 MADs above the median for every 5-min period (Gunaydin et al., 2014; Muir et al., 2018). The transient frequency for each 5-min bin was then averaged within groups. To restrict detection to absolute peaks, a minimum of 0.3 seconds between peaks and prominence of 1 MAD above the surrounding signal was required (MATLAB).

Blood glucose mobilization in response to behavioral stress was assessed as a measure of sympathetic activation (Bialik et al., 1988). Contour Next EZ glucometers (Bayer, Parsippany, NJ) were used to generate duplicate readings for each time point that were then averaged. For plasma analysis of corticosterone, the primary product of the rat hypothalamic-pituitary-adrenal axis (Herman et al., 2016; Ghosal et al., 2017b), blood samples were centrifuged at 3000 X g for 15 minutes at 4° C with plasma stored at −20° C until ELISA. Plasma corticosterone was measured with an ENZO Corticosterone ELISA (ENZO Life Sciences, Farmingdale, NY) with an intra-assay coefficient of variation of 8.4% and an inter-assay coefficient of variation of 8.2% (Bekhbat et al., 2018; Dearing et al., 2021). Samples were analyzed in duplicate with all time points included in the same assay.

### Food reward

To measure IL^CaMKIIα^ neural activity during reward acquisition, animals were fed chocolate. For 2 days at both lights on and off, animals were acclimated to being moved to an adjacent behavioral testing room with a chocolate chip (Ghiradelli, semi-sweet chocolate morsels) placed in the home cage. On the testing day, rats were connected to the fiber photometry recording system and placed in their home cage for 3 min prior to adding the chocolate chip to the cage. Animals taking > 30 min to consume the chocolate were excluded from analysis. Neural activity during chocolate interactions was analyzed as described above for object and social interactions.

### Tissue collection

At the conclusion of experiments, rats were given an overdose of sodium pentobarbital and perfused transcardially with 0.9% saline followed by 4.0% paraformaldehyde in 0.1 M PBS. Brains were removed and post-fixed in 4.0% paraformaldehyde for 24 h at room temperature, followed by storage in 30% sucrose in PBS at 4 °C. Coronal sections were made on a freezing microtome at 30 µm thickness and then stored in cryoprotectant solution at − 20 °C prior to localizing viral injections and optic cannula placement.

### Statistical analysis

Data are expressed as mean ± standard error of the mean. Data were analyzed using Prism 8 (GraphPad, San Diego, CA), with statistical significance set at *p <* 0.05 for rejection of null hypotheses. Weights by treatment (CVS) and day were analysed using repeated-measure 2-way analysis of variance (ANOVA), followed by Fisher’s post hoc if significant main or interaction effects were present. Object, social, and reward interactions were analyzed via unpaired t-tests between stress or interaction condition. Peak neural activity preceding social interaction, as well as glucose and corticosterone across restraint days were analyzed using mixed-effects analysis with stress and time (repeated) as factors, followed by Fisher’s post hoc if significant main or interaction effects were present. Transient frequency during restraint was analyzed using mixed-effects analysis with home cage/restraint (repeated) and CVS as factors followed by Sidak’s post hoc if significant main or interaction effects were present. Male body weight change and mean chocolate Z-score comparisons across males, females, and OVX females were analyzed by 1-way ANOVA, followed by Fisher’s LSD post hoc if significant main or interaction effects were present.

## Results

### Design and validation

AAV viral injections were targeted to the IL (**Figure 1A**) for expression of GCaMP6s under CaMKIIα promoter regulation (IL^CaMKIIα^). Fiber optic cannulas were targeted to the IL to allow excitation and collection of GCaMP6s fluorescence (Gunaydin et al., 2014; Muir et al., 2018). All experiments (male, female, and OVX) underwent the same experimental design (**Figure 1B**). Only animals with cannula placement verified to be within or immediately (≤ 0.2 mm) dorsal to the IL were included in analyses (**Figure 1C**). In all experiments, No CVS day 13 weight was higher than day 1. This effect was not present in any CVS group and male and OVX females had lower body weight than No CVS controls (Males: repeated-measures 2-way ANOVA: Time x Stress F_(1, 19)_ = 10.66, *p <* 0.01; Females: repeated-measures 2-way ANOVA: Time x Stress F_(1, 18)_ = 8.15, *p <* 0.05; OVX Females: repeated-measures 2-way ANOVA: Time x Stress F_(1, 20)_ = 209.2, *p <* 0.0001) (**Figure 1D**). As previously reported (Chen and Heiman, 2001; Fang et al., 2015), OVX females had higher body weight before CVS than cycling females (n = 20-22/group, unpaired t-test: Female vs OVX Female weight t_(40)_ = 18.61, *p <* 0.0001). Intact female estrous phase was determined following each testing session (**Table 1**); however, sample sizes did provide the statistical power required to analyze phase-dependent effects on neural activity.

**Table 1:**
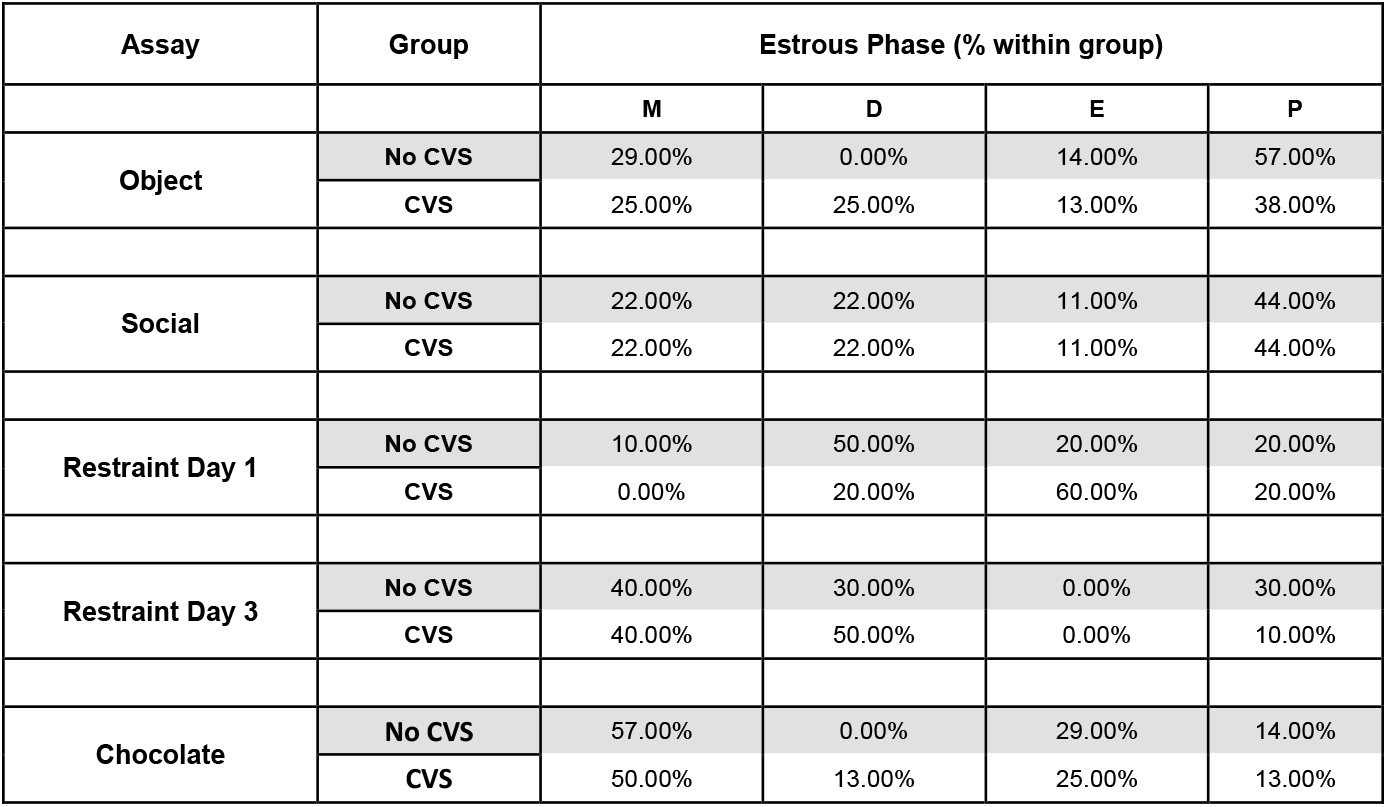
Distribution of estrous cycle phase during testing. M: metestrus, D: diestrus: E: estrus, P: proestrus.

### Object interactions

To determine how IL^CaMKIIα^ neurons encode interactions with a novel object, an empty enclosure was placed in the arena (**Figure 2A**). During the first object interaction (**Figure 2B**), CVS males had a greater GCaMP signal than No CVS males (n = 8-10/group, unpaired t-test: No CVS vs CVS t_(16)_ = 2.483, *p <* 0.05; **Figure 2C**). While all No CVS groups had increased activity (**Figure 2D**) during subsequent object interactions (Random vs Object unpaired t-test: Males: n = 8, t_(14)_ = 4.75, *p <* 0.001; Females: n = 7, t_(12)_ = 5.8, *p <* 0.0001; OVX: n = 5, t_(8)_ = 3.1, *p <* 0.05; **Figure 2E**), there was no effect of CVS in any experiment to alter the encoding of a familiar object. Although, activity during the initiation (first 0.5 sec) of interactions trended upward in intact females (Females unpaired t-test: n = 7-8/group, t_(13)_ = 2.12, p = 0.054; **Figure 2F**). Overall, chronic stress increases male IL^CaMKIIα^ neuronal responses to novelty, an effect not present in females regardless of ovarian status.

**Figure 2:**
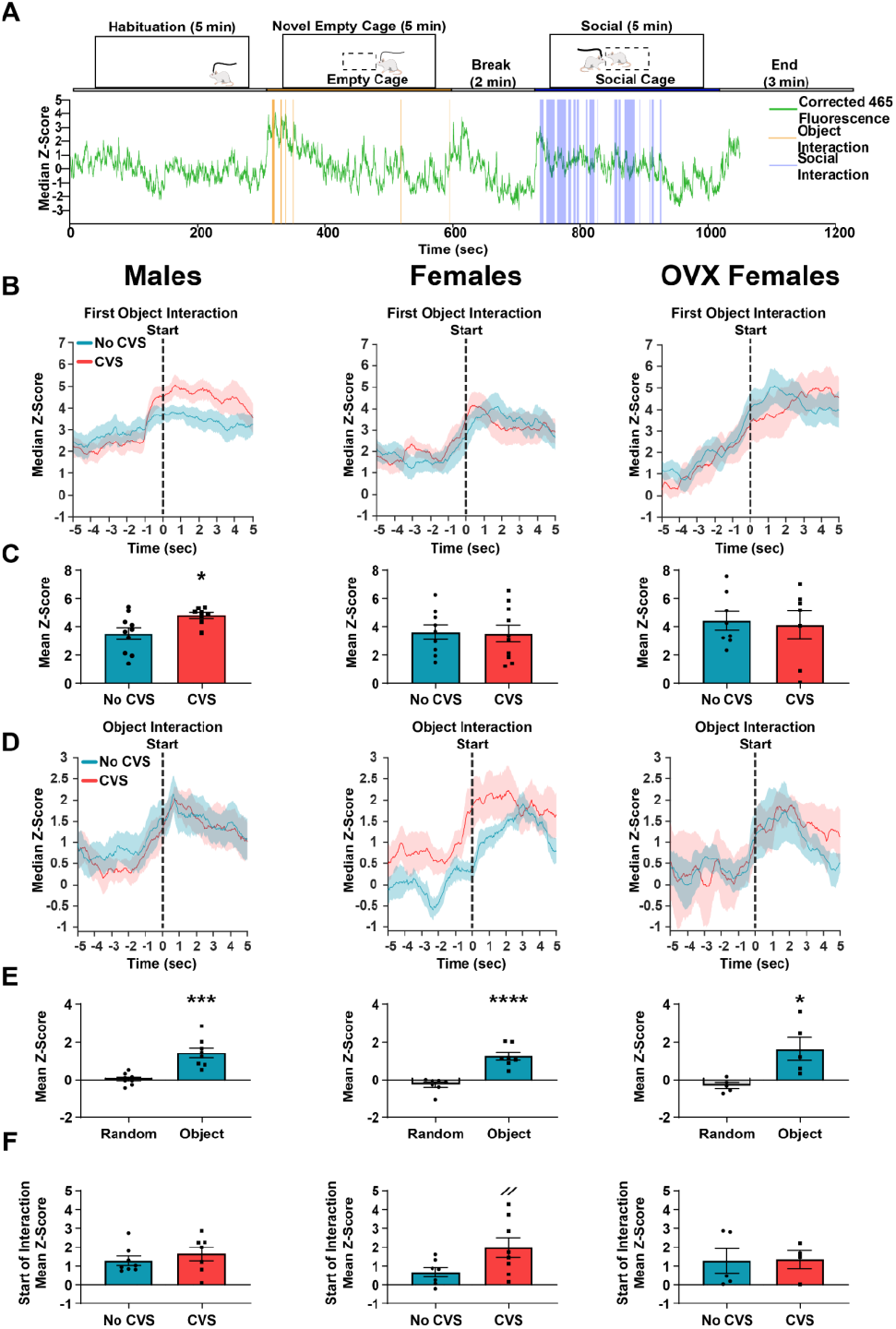
IL^CaMKIIα^ neurons responded to object interactions. (**A**) Schematic of experimental approach with a representative median-corrected 465 nm signal trace (green). (**B**) Group averaged z scores aligned to the first object interaction, the dashed line represents interaction start; OVX: ovariectomized. (**C**) CVS exposure increased neural activity during the first object interaction in males only. (**D**) Group averaged z scores aligned to subsequent object interactions. (**E**) All No CVS groups had higher mean Z-scores during object interactions compared to randomly selected non-specific behaviors. (**F**) CVS trended (p = 0.054) toward increased signal at the start of object interactions only in cycling females. Comparisons between treatments: ^//^ *p <* 0.06, * *p <* 0.05, *** *p <* 0.001, **** *p <* 0.0001.

### Social Interactions

Following object interaction, a novel conspecific was added to the arena to examine IL representation of social interaction. There were no CVS effects in any experiment on the GCaMP signal during the first social interaction (data not shown). For subsequent social interactions (**Figure 3A**), No CVS males, females, and OVX females all had higher mean IL^CaMKIIα^ neural activity during social interactions than random intervals (Random vs Social unpaired t-test: Males: n = 8, t_(14)_ = 5.3, *p <* 0.0001; Females: n = 9, t_(16)_ = 5.28, *p <* 0.0001; OVX Females: n = 7, t_(12)_ = 8.05, *p <* 0.0001; **Figure 3B**). Further, CVS reduced the mean GCaMP signal during social interaction in males (**Figure 3C**). Females were resistant to CVS effects on social representation, which was dependent on ovarian hormones (No CVS vs CVS unpaired t-tests: Males: n = 8/group, t_(14)_ = 2.19, *p <* 0.05; OVX Females: n = 7/group, t_(12)_ = 3.09, *p <* 0.01). Neural activity immediately preceding social interaction was assessed to examine the neural basis of social motivation. In males, the IL^CaMKIIα^ GCaMP signal increased prior to the start of social interaction (Males mixed-effects model: Time F_(2.648, 32.43)_ = 5.2, *p <* 0.01; **Figure 3D**), which CVS reduced (*p <* 0.05). In contrast, there were no time-dependent changes in either intact or OVX females. Taken together, chronic stress reduced the prefrontal representation of social interaction in males and OVX females without altering responses in intact females. Further, male IL activity increased prior to social interaction, an effect that was reduced by chronic stress.

**Figure 3:**
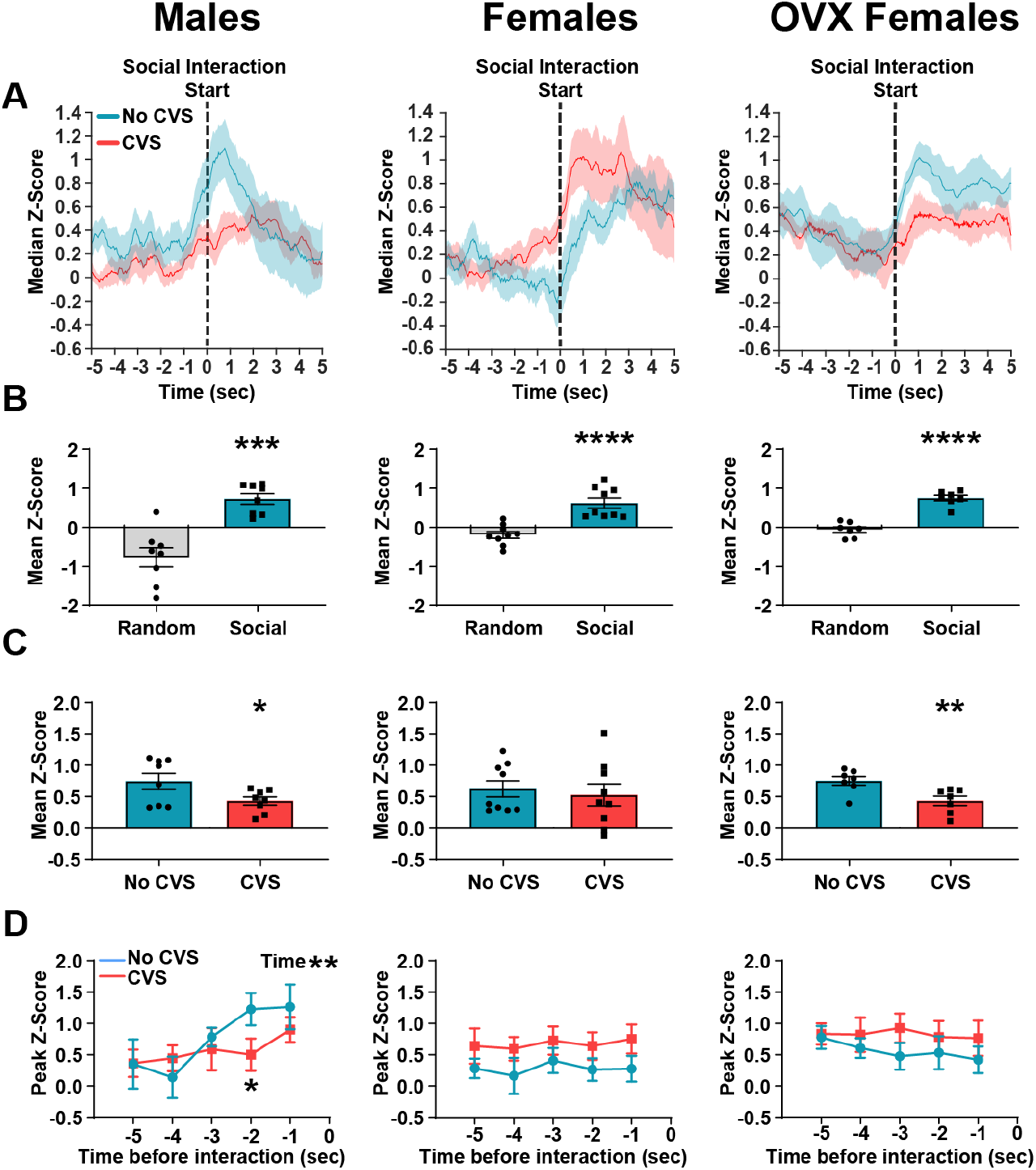
CVS reduced IL^CaMKIIα^ neural activity during social interactions in males and OVX females. (**A**) Group averaged z scores aligned to social interactions, the dashed line represents interaction start; OVX: ovariectomized. (**B**) All No CVS groups had higher mean Z-scores during object interactions compared to randomly selected non-interaction periods. (**C**) CVS exposure reduced mean z score during social interactions in males and OVX females, without effecting cycling females. (**D**) Male GCaMP signal increased prior to social interactions, which was reduced by CVS. Females and OVX females showed no change in neural activity preceding interactions. Comparisons between treatments: * *p <* 0.05, ** *p <* 0.01, *** *p <* 0.001, **** *p <* 0.0001.

### Restraint stress

Restraint was used to examine IL^CaMKIIα^ neuronal responses to an acute novel stressor. Restraint stress was then repeated (3 total days) to query neural encoding after repeated stress. After each restraint session, blood was sampled to measure glucose and corticosterone as indices of the physiological stress response. In males, CVS increased corticosterone after restraint on the second and third days (mixed-effects model: Stress F_(1, 27)_ = 11.99, *p <* 0.01; **Figure 4B**). In cycling females there was an interaction between day and CVS (repeated-measures 2-way ANOVA: Day x Stress F _(2, 32)_ = 3.57, *p* < 0.05) where CVS increased corticosterone on day 1 (*p <* 0.05). Similarly, OVX females had a day and CVS interaction (mixed-effects model: Day x Stress F_(2, 39)_ = 4.66, *p <* 0.05; **Figure 4B**), with CVS increasing corticosterone on day 1 (*p <* 0.01). Additionally, OVX No CVS females had higher corticosterone on day 2 than day 1 (*p <* 0.05). Consistent with previous research (Viau and Meaney, 1991), cycling No CVS females had higher corticosterone stress responses than No CVS OVX females, (unpaired t-test Female vs OVX Female: n = 9-11/group, t_(18)_ = 2.76, *p <* 0.05). Altogether, corticosterone data indicate that, relative to No CVS controls, CVS caused a progressive HPA axis sensitization in males contrasted by a transient facilitation in both female experiments.

**Figure 4:**
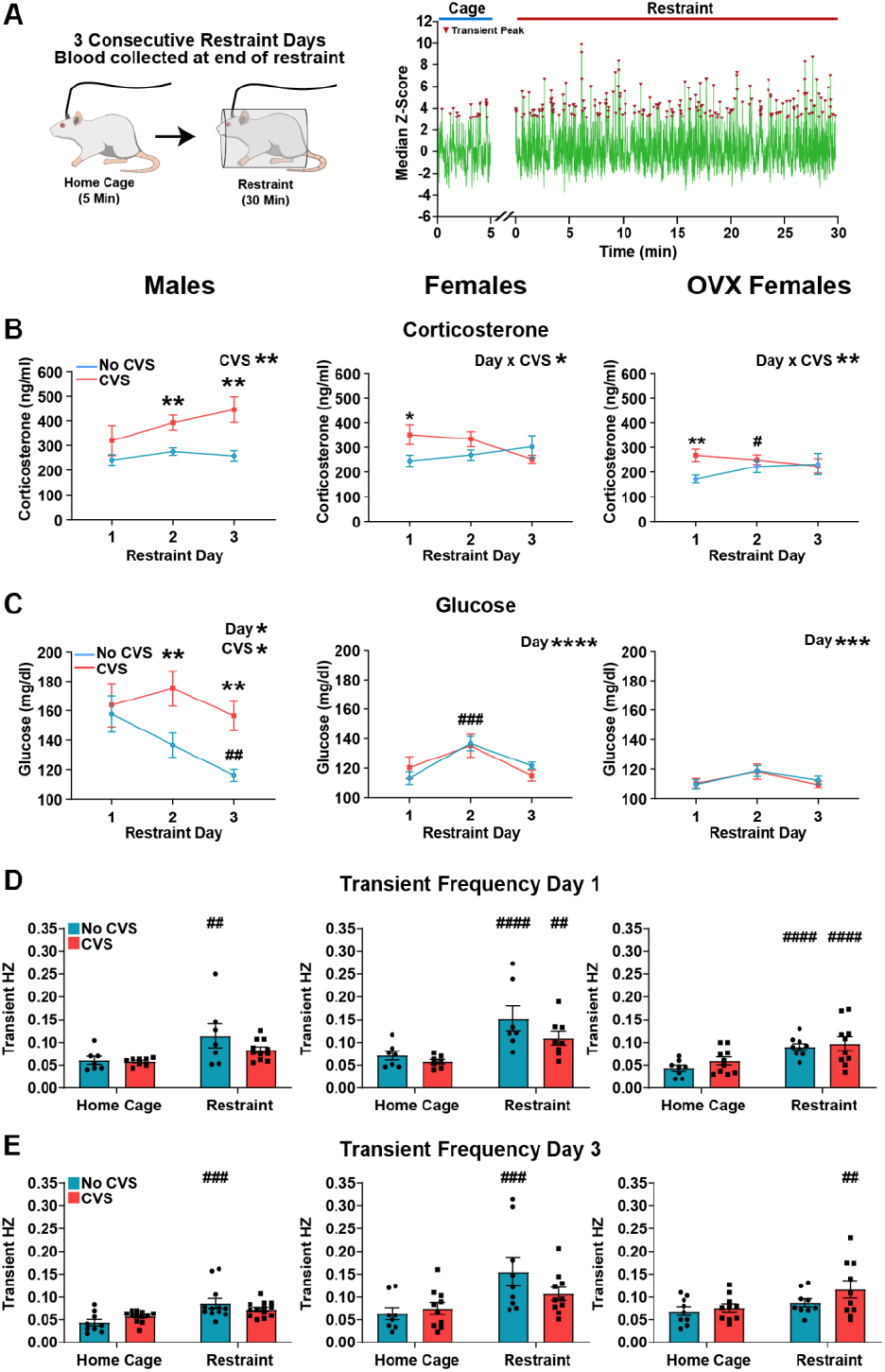
IL^CaMKIIα^ neurons responded to acute and repeated stress. (**A**) Left: Schematic of restraint approach. Right: Representative transient peak quantification during home cage and restraint. (**B**) CVS increased corticosterone responses in males on days 2 and 3, while females and OVX females had day 1 increases; OVX: ovariectomized. (**C**) CVS exposure increased male glucose responses on days 2 and 3. There was no CVS effect in females or OVX females. (**D**) Acute novel restraint. In males, only No CVS animals had higher transient frequency during restraint than home cage. In intact and OVX females, both No CVS and CVS rats had higher transient frequency during restraint. (**E**) Day 3 repeated restraint. No CVS males had higher transient frequency during restraint than home cage. Intact No CVS females had higher transient frequency during restraint, while OVX CVS females had higher transient frequency. Treatment effects: * *p <* 0.05, ** *p <* 0.01. Within-treatment time comparisons: ^#^ *p <* 0.05, ^##^ *p <* 0.01, ^###^ *p <* 0.001, ^####^ *p <* 0.0001.

There were main effects of CVS and day on male glucose responses with CVS-exposed animals having higher glucose mobilization on days 2 and 3 than No CVS controls (mixed-effects model: Day F_(1.588, 39.69)_ = 4.912, *p <* 0.05; Stress F_(1, 28)_ = 5.13, *p <* 0.05; **Figure 4C**). Also, No CVS males had lower glucose on day 3 than day 1 (*p <* 0.01), a habituation that was prevented by CVS. In both intact and OVX females there was a main effect of restraint day (mixed-effects model Females: Day F_(1.882,31.05)_ = 13.66, *p <* 0.0001; OVX Females: Day F_(1.887,33.97)_ = 4.43, *p <* 0.05) where cycling No CVS females had higher glucose on day 2 than day 1 (*p <* 0.05). Thus, chronic stress prevented habituation of sympathetic glucose mobilization in males without altering glucose responses in intact or OVX females.

Neural activity was measured as IL^CaMKIIα^ transient peak frequency using rolling 1-minute medians and MAD Z-scores. Recordings occurred in the home cage and immediately following in restraint. On day 1 in males, No CVS animals had higher transient frequency during restraint compared to home cage, while CVS animals did not (mixed-effects model: Restraint F_(1, 13)_ = 19.50, *p <* 0.001 Sidak’s: No CVS: *p <* 0.01; **Figure 4D**). In contrast, both female and OVX female stress conditions had higher transient frequency during restraint than home cage (mixed-effects model: Females Restraint F_(1, 15)_ = 36.35, *p <* 0.0001, Sidak’s: No CVS: *p <* 0.001, CVS: *p <* 0.01; OVX females Restraint F_(1, 16)_ = 70.07, *p <* 0.0001, Sidak’s: No CVS: *p <* 0.0001, CVS: *p <* 0.0001).

On the third day of repeated restraint, male responses were similar to day 1 with a main effect of restraint that was significant only in the No CVS group (mixed-effects model: Males Restraint F_(1, 19)_ = 20.16, No CVS *p <* 0.001; **Figure 4E**). Cycling females had population transient frequencies similar to males with an effect of restraint in the No CVS group only (mixed-effects model: Restraint F_(1, 16)_ = 24.85, *p <* 0.001, No CVS *p <* 0.001). However, OVX females had an effect of restraint where CVS animals had higher transient frequency (mixed-effects model: Restraint F_(1, 17)_ = 14.56, CVS *p <* 0.01). Altogether, the data suggest that stress increased IL^CaMKIIα^ calcium transients only in No CVS males. Regardless of stress condition, both intact and OVX females had increased calcium transients during novel restraint, followed by diverging condition-specific responses on day 3.

To assess associations between IL^CaMKIIα^ neural activity during stress and endocrine responses, Pearson’s correlations were calculated between transient frequency and glucose or corticosterone (**Table 2**). Only in males, No CVS IL^CaMKIIα^ population activity positively correlated with glucose (day 2; *p <* 0.05; day 3; *p <* 0.01) and corticosterone (day 3; *p <* 0.001) during repeated restraint. In contrast, acute novel restraint (day 1; *p <* 0.05) led to a negative correlation between male CVS neural activity and stress-evoked glucose. There were no significant correlations in any female condition between IL^CaMKIIα^ transient frequency during restraint and glucose or corticosterone.

**Table 2:** Pearson’s coefficients of correlation between mean daily IL^CaMKIIα^ calcium transients during restraint and post-stress glucose and corticosterone.

|  |  | Males |  |  |  | Females |  |  |  | OVX Females |  |  |  |
| --- | --- | --- | --- | --- | --- | --- | --- | --- | --- | --- | --- | --- | --- |
|  |  | Glucose |  | Corticosterone |  | Glucose |  | Corticosterone |  | Glucose |  | Corticosterone |  |
| Day | Group | R | P | r | p | R | p | r | p | r | p | R | p |
| 1 | No CVS | 0.60 | 0.11 | 0.45 | 0.26 | -0.60 | 0.12 | 0.30 | 0.46 | -0.21 | 0.59 | 0.24 | 0.54 |
| 1 | CVS | -0.63 | 0.04 | -0.56 | 0.09 | -0.13 | 0.75 | -0.42 | 0.30 | -0.49 | 0.15 | -0.07 | 0.85 |
| 2 | No CVS | 0.66 | 0.04 | 0.48 | 0.16 | -0.03 | 0.94 | 0.14 | 0.72 | -0.25 | 0.48 | 0.54 | 0.11 |
| 2 | CVS | 0.49 | 0.07 | -0.14 | 0.64 | 0.25 | 0.49 | -0.39 | 0.26 | 0.53 | 0.10 | -0.04 | 0.91 |
| 3 | No CVS | 0.71 | 0.01 | 0.84 | 0.001 | 0.58 | 0.10 | -0.22 | 0.58 | 0.07 | 0.86 | 0.39 | 0.30 |
| 3 | CVS | 0.32 | 0.29 | 0.26 | 0.39 | -0.19 | 0.65 | -0.02 | 0.97 | -0.21 | 0.56 | -0.28 | 0.43 |

### Food reward

Chocolate chips were given as a palatable food to assess IL^CaMKIIα^ neural responses to reward. To reduce novelty, rats were acclimated to chocolate chips in the home cage for 2 days prior to recording. Controlling for potential effects of energy balance, Experiment 1 had an additional group of males that were food restricted to match the body weight change of CVS males (**Figure 5A**). Food-restricted males had 2-week body weight changes similar to CVS-exposed rats, with both groups reduced compared to No CVS males (1-way ANOVA: n = 7-14/group, F_(2,26)_ = 11.07, *p <* 0.001, Tukey post hoc; **Figure 5B**). Photometry data were centered on grabbing events when the chocolate reward was acquired (**Figure. 5C**). No CVS rats in all experiments had higher neural activity during the first half-second of grabbing chocolate than randomly selected periods (unpaired t-test random vs chocolate: Males: n = 7/group, t_(12)_ = 3.55, *p <* 0.01; Females: n = 9/group, t_(16)_ = 5.154, *p <* 0.0001; OVX Females: n = 6/group, t_(10)_ = 2.664, *p <* 0.05: **Figure 5D**). Further, intact No CVS females had higher IL^CaMKIIα^ GCaMP signal at chocolate acquisition than No CVS males or OVX females (1-way ANOVA: n = 6-9, F_(2,19)_ = 3.822, *p <* 0.05). While CVS males had increased GCaMP fluorescence with chocolate acquisition, food-restricted males did as well (1-way ANOVA: n = 7-11, F_(2,20)_ = 3.466, p = 0.05; **Figure 5E**). There were no CVS effects in either female or OVX female experiments. Overall, CVS increased IL^CaMKIIα^ neural activity during food reward in males, an effect that was accounted for by reduced body weight gain. Cycling No CVS females had ovarian hormone-dependent elevations in neural activity, but CVS did not alter reward processing in either female experiment.

**Figure 5:**
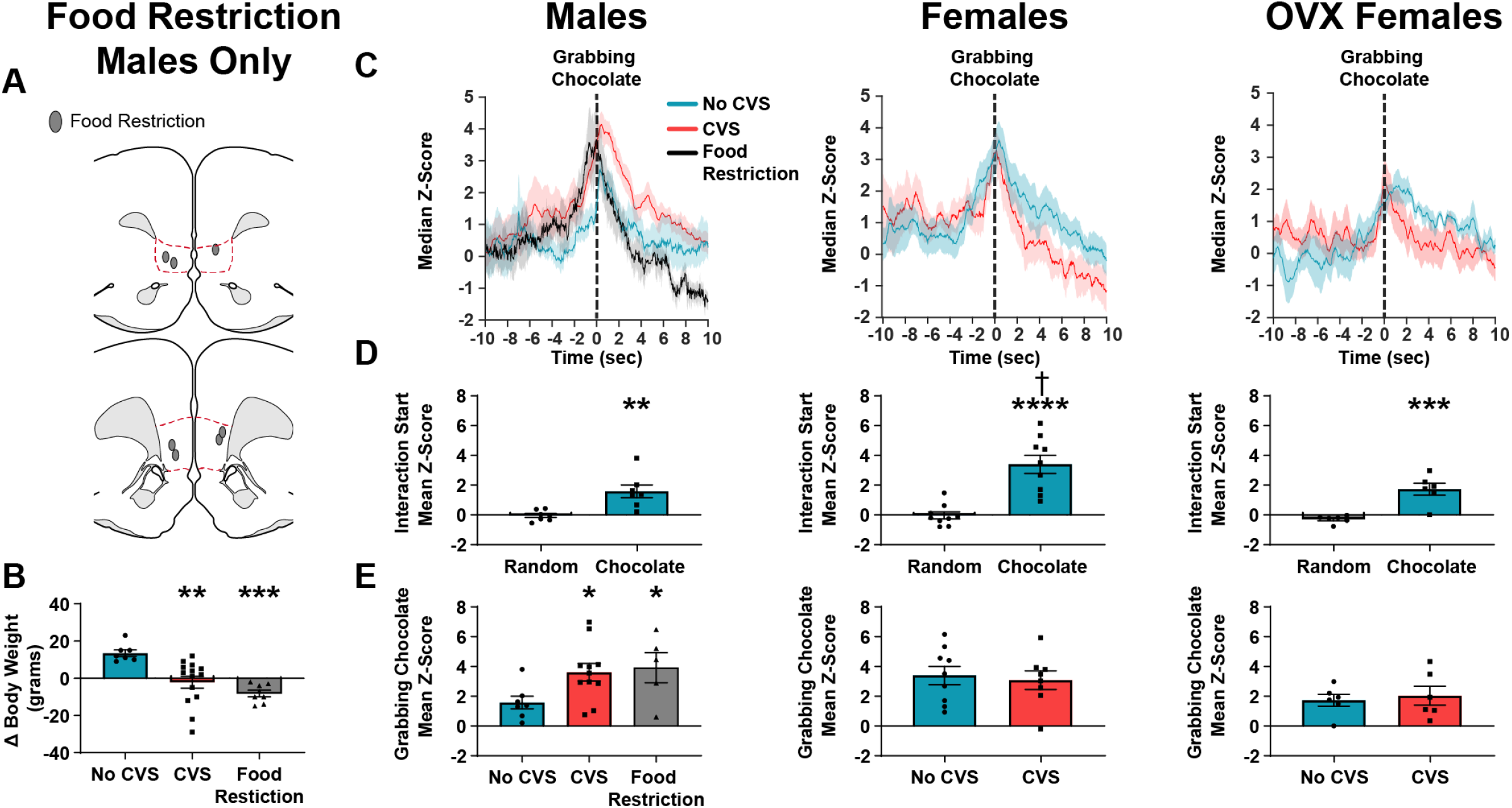
CVS and food restriction increased male IL^CaMKIIα^ neuronal responses to food reward. (**A**) Mapped fiber-optic cannula positions within the IL (red outline). (**B**) Food restriction reduced body weight gain similar to CVS. (**C**) Group averaged z scores were aligned to chocolate acquisition, dashed line represents rat grabbing chocolate; OVX: ovariectomized. (**D**) All groups had higher mean z scores during the first half-second of grabbing the chocolate compared to randomly selected non-interaction periods. Cycling females had greater IL pyramidal cell responses than males or OVX females. (**E**) CVS exposure and food restriction increased GCaMP signaling in males, with no effect in cycling or OVX females. Treatment effects: * *p <* 0.05, ** *p <* 0.01, *** *p <* 0.001, **** *p <* 0.0001. Sex effects: ^†^ *p <* 0.05.

## Discussion

The current studies utilized fluorescent calcium imaging of genetically-defined IL^CaMKIIα^ neurons in male and female rats. Recordings of neural activity occurred during real-time behavior as animals engaged affective stimuli. Both negatively-valanced stimuli (e.g. stress) and positively-valanced paradigms (including food reward) assessed cortical processing of emotion-relevant behaviors. The collective results indicate that both male and female IL^CaMKIIα^ neurons increase activity during novel, social, and reward interactions compared to non-specific behaviors in the testing environment. Further, comparisons across sex in No CVS animals found similar neural responses to novelty, social interaction, and stress in males and females. However, cycling females had increased responses to food reward, which was prevented by ovariectomy. Additionally, only males had time-dependent neural activity during social approach.

Models of chronic heterotypic stress (e.g. CVS, chronic mild stress, chronic unpredictable stress) are widely used to study the neural, behavioral, and physiological impacts of chronic stress. Numerous studies in male and female rodents report that CVS and related paradigms promote novelty avoidance (Muir et al., 2020; Pace et al., 2020b), reduced sociability (Franceschelli et al., 2014; Muir et al., 2020), and anhedonia (Willner, 2005; Hersey et al., 2022), core features of affective disorders. Here, we examined the consequences of CVS on *in vivo* IL^CaMKIIα^ neural activity during behavior. Our results indicate that chronic stress-induced outcomes are dependent on both sex and behavioral context. Specifically, CVS increased male GCaMP signaling during interactions with a novel, but not familiar, object. Moreover, male IL^CaMKIIα^ neural activity increased during acquisition of chocolate food reward, an effect accounted for by metabolic status. In contrast, CVS reduced male responses during social approach and interaction. CVS in males also prevented stress-induced increases in IL^CaMKIIα^ transient frequency during novel and repeated restraint, as well as increasing endocrine stress responses over the course of repeated restraint.

Compared to No CVS controls, chronically-stressed cycling female rats did not have significant changes in IL^CaMKIIα^ calcium signaling during behavior. Sex differences in the consequences of CVS were partially accounted for by ovarian hormones. Similar to males, CVS reduced social interaction encoding in OVX females. Thus, ovarian factors prevented the effects of CVS on IL^CaMKIIα^ representation of social interaction. However, prevention of other CVS effects was independent of ovarian status. For instance, neither cycling nor OVX females were susceptible to CVS effects on social approach or novel object encoding. During repeated restraint, cycling and OVX CVS females also had similar patterns of physiological stress responses relative to respective No CVS controls. In addition, novel restraint after CVS increased IL^CaMKIIα^ calcium transients in both female conditions. Altogether, the results point toward multiple sex differences in chronic stress-induced outcomes that are independent of the activational effects of ovarian hormones. Although male metabolic state accounted for CVS effects on food reward and ovarian status explained sex differences in social encoding after CVS, the biological bases underlying broader sex differences in novelty and stress responses remain to be determined. Ovariectomy prevents the activational effects of ovarian hormones, yet the potential remains for developmental effects as well as hormone-independent chromosomal effects (Sheng et al., 2021). Further, gonadal hormone receptor modulation by neurosteroids (Wei et al., 2014; Brann et al., 2021) and/or ligand-independent signaling (Pak et al., 2006; Pinceti et al., 2015) may also contribute. Ultimately, multiple interacting factors may account for sex differences in prefrontal activity after chronic stress.

In addition to the sex-specific impacts of chronic stress on prefrontal cortex activity, male and female IL^CaMKIIα^ neurons also divergently modulate behavioral and physiological responses to stress (Wallace and Myers, 2021). Glutamate output from the male IL is necessary for chronic stress-induced avoidance behaviors (Pace et al., 2020b) and constrains physiological stress responses (Myers et al., 2017; Schaeuble et al., 2019). Furthermore, optogenetic stimulation of male IL^CaMKIIα^ neurons increases place preference and social motivation, in addition to reducing sympathetic and HPA axis responses to stress (Wallace et al., 2021). In contrast, female IL^CaMKIIα^ stimulation increases sympathetic stress responding without modulating social or motivational behaviors. Combined with the results of the current experiments, these findings point toward an interesting female divergence of behavioral regulation and behavior processing where female IL^CaMKIIa^ neurons respond to social interaction without modulating social motivation. Additionally, the differential impact of male and female IL activity on physiological stress responding may contribute to the male-specific correlations between calcium transient frequency and endocrine responses during restraint.

Histological studies have examined the impact of chronic stress on vmPFC morphology with multiple reports of male pyramidal cell dendritic hypotrophy (Cerqueira et al., 2005; Goldwater et al., 2009; Shansky et al., 2009; Luczynski et al., 2015; Czéh et al., 2018). However, changes in male local inhibitory networks are mixed. Chronic stress increases dendritic arborization of GAD67-positive interneurons while reducing the number of GABAergic interneurons (Gilabert-Juan et al., 2013). Additional evidence for stress-induced male vmPFC reorganization comes from studies identifying both elevated (Shepard et al., 2016) and reduced GAD67 mRNA (Ghosal et al., 2020). The functional consequences of altered interneuronal morphology and gene expression have been queried with *ex vivo* slice physiology. To date, this approach has also yielded reports of both increased (McKlveen et al., 2016) and decreased (Czéh et al., 2018) inhibitory currents on male IL pyramidal cells after CVS. Accordingly, hypotheses have developed of vmPFC hyper- or hypo-inhibition after chronic stress (Ghosal et al., 2017a; Fogaça and Duman, 2019; McKlveen et al., 2019; Page and Coutellier, 2019). The current studies advance this framework by showing that, during *in vivo* behavior, chronic stress dynamically shifts male excitatory/inhibitory balance in a context-dependent manner. In this case, responses to novelty and reward are increased while social and stress responses are decreased. Ultimately, both hypo- and hyper-function of the male vmPFC may produce context-inappropriate behaviors and underlie multiple aspects of stress pathology.

Regarding females, no studies to our knowledge have investigated chronic heterotypic stress effects on IL-specific pyramidal cell structure or function. Investigations of pyramidal morphology in other PFC regions have yielded a variety of outcomes, likely as a function of stress paradigm. For instance, the prelimbic cortex has similar reductions in pyramidal spine density in males and females after CVS but fewer impairments after chronic restraint stress (Anderson et al., 2019). However, shorter durations of chronic restraint stress produce estradiol-dependent female dendritic proliferation in prelimbic cortex (Garrett and Wellman, 2009). Interestingly, chronic restraint stress affects IL pyramidal cells in a circuit-specific and hormone-dependent manner. Here, OVX females have general increases in IL spine density after chronic restraint that are prevented by estradiol replacement (Shansky et al., 2010). However, amygdala-projecting IL neurons have increased spine density following chronic restraint regardless of hormonal status (Shansky et al., 2010). Female PFC interneurons are also susceptible to structural changes following prolonged stress. In a CVS-like paradigm, females, but not males, have increased parvalbumin mRNA in the PFC (Shepard et al., 2016). More specific to the IL, chronic stress also increases the density of female parvalbumin-positive interneurons (Shepard et al., 2016). It is unclear how increased female IL parvalbumin cells might impact the function of local circuits. However, broad PFC slice recordings after chronic restraint indicate that females do not have the reduced glutamatergic transmission described in males (Yuen et al., 2012; Wei et al., 2014). This sex difference is accounted for by estradiol signaling at estrogen receptors in the PFC; although, the source of estradiol remains to be determined as the signaling is ovary-independent and reliant on aromatase (Wei et al., 2014). In our studies, females did not exhibit CVS-induced changes in IL^CaMKIIα^ activity. Ovarian hormones prevented the impact of CVS on social encoding but the absence of CVS effects in other contexts was ovary independent. Given that female IL^CaMKIIα^ cells facilitate sympathetic stress responses (Wallace and Myers, 2021), it is unclear whether this resistance to CVS is a beneficial or deleterious outcome.

While the current studies significantly advance understanding of *in vivo* neural activity in chronically-stressed males and females, there are multiple methodological limitations worth noting. One limitation of bulk population recordings is the inability to identify neural ensembles that may be stimulus- or behavior-specific. The degree of cellular diversity in response characteristics across stimuli remains unknown, as does the potential overlap of cellular populations encoding social, stress, and reward stimuli. Further, heterogeneity may arise from output-defined ensembles. Thus, specific cortical projections may be differentially sensitive to chronic stress in a sex-dependent manner. Another interpretive consideration is the relatively slow kinetics of the GCaMP fluorescent indicator. Previous studies have demonstrated that photometry recordings are reflective of summated neural activity, yet the kinetics of GCaMP do not permit action potential-level resolution (London et al., 2018). Furthermore, the photometry approach is a normalized measure of relative changes in calcium signaling and does not quantify absolute neural activity. Thus, the required baseline corrections for event-centered analyses may limit detection of sex or stress effects on total neural activity.

In summary, the current experiments provide *in vivo* recordings of IL pyramidal cell activity in males and females. Response characteristics were similar across sexes with the exceptions of social approach and food reward. However, chronic stress exposure predominately altered male IL^CaMKIIα^ neural activity. Notably, CVS did not globally increase or decrease male neural activity, instead chronic stress altered activity depending on behavioral context. Thus, the effects of chronic stress on male IL^CaMKIIα^ population activity are highly context dependent. Although male metabolic and female hormonal status partially explain sex differences in responses to chronic stress, other outcomes arise from different mechanisms. Ultimately, the data shed light on the importance of behavioral context and sex for understanding cortical processing of affective stimuli after chronic stress and suggest that these factors require consideration in the effort to understand and treat affective illness.

## Acknowledgements

This work was supported by the College of Veterinary Medicine and Biomedical Sciences Research Council and NIH grant HL150559 to Brent Myers. Addgene viral prep #107790-AAV9 was made available by a gift from James M. Wilson.

